# Radiolabelled Bacterial Metallophores as Targeted PET Imaging Contrast Agents for Accurate Identification of Bacteria and Outer Membrane Vesicles *in vivo*

**DOI:** 10.1101/2020.08.06.240119

**Authors:** Nabil A. Siddiqui, Hailey A. Houson, Shindu C. Thomas, Jose R. Blanco, Robert E. O’Donnell, Daniel J. Hassett, Suzanne E. Lapi, Nalinikanth Kotagiri

## Abstract

Modern technologies such as 16s DNA sequencing capable of identifying microbes and provide taxonomic resolution at species and strain-specific levels is destined to be transformative^1^. Likewise, there is an emerging need to accurately identify both infectious and non-infectious microbes non-invasively in the body at the genus and species level to guide diagnosis and treatment strategies. Here, we report development of radiometal-labelled bacterial chelators, knowns as metallophores that allow non-invasive and selective imaging of bacteria and bacterial products *in vivo*. We show that these novel contrast agents are able to identify *E. coli* with strain level specificity and other bacteria, such as *K. pneumoniae*, based on expression of distinct cognate transporters on the bacterial surface. The probe is also capable of tracking probiotic, engineered bacteria and bacterial products, outer membrane vesicles (OMVs), in unique niches such as tumours. Moreover, we report that this novel targeted imaging approach has impactful applicability in monitoring antibiotic treatment outcomes in patients with pulmonary infections, thereby providing the ability to optimize individualized therapeutic approaches. Compared to traditional techniques used to manufacture probes, this strategy simplifies the process considerably by combining the function of metal attachment and cell recognition into a single molecule. Thus, we anticipate that these probes will be widely used in both clinical and investigative settings in living systems for non-invasive imaging of infectious and non-infectious organisms.

---

Non-invasive imaging of bacteria, particularly “live” bacteria in the body is in its infancy and a growing critical need, especially during infection. Medically, the precise identification and localization of such organisms in the human body is of vital importance. An effective strategy would be to eliminate diagnostic uncertainty so that selection of antibiotic treatment is often empiric^2^. Ideally, such approaches should also be able to determine their distribution throughout the body, and distinguish pathogenic from commensal species to minimize indiscriminate use of broad-spectrum antibiotics thereby adopting a more focused treatment strategy targeting specific pathogens. An application of equal importance is the precise mapping of resident microbiota, tracking of engineered bacteria and their products such as virulence factor-harbouring outer membrane vesicles (OMVs) for therapeutic applications. The role of microbiota in human disease is increasingly being recognized to play a major role in aetiology and treatment modifications^3^. The emergence of next-generation sequencing techniques has allowed extensive characterization of the microbiome in various pathologic conditions, from neurologic and endocrine, to cancer^4-6^. However, there is an urgent need for non-invasive tools to detect and identify bacteria using endogenous markers. Bacteria are also commonly used for desirable, biotechnological applications due to the ease of genetic manipulation and heterologous protein expression. Several broad categories of bacteria-based approaches have been explored in preclinical research focusing on a wide range of diseases such as cancer, diabetes, obesity, hypertension, infection and inflammatory dysfunction^7-10^. However, functional stability is important for biomedical applications of engineered bacteria. Development of convenient and realistic *in vivo* testing environments will be important for achieving improved accuracy. Such monitoring would provide information regarding presence, location, quantity, proliferation, survival and status of the current microbiome, engineered bacteria and OMVs.

Numerous molecular imaging techniques such as ultrasound (US), computed tomography (CT), magnetic resonance imaging (MRI), single-photon emission CT (SPECT), and positron emission tomography (PET) have been developed for preclinical and clinical research in the last three decades. However, current clinical probes cannot reliably distinguish between bacteria from mammalian cells *in vivo*^11^. Thus, a comprehensive understanding of bacterial physiology and genetics is required to develop probes for targeted imaging. As bacteria are evolutionarily and phylogenetically distinct from mammalian cells, fundamental differences in metabolism and cellular structures can be leveraged to develop bacteria-specific imaging agents. Some of the recent approaches that have focused on targeting metabolic pathways and proteins that are only present in bacteria include – ^11^C-para-aminobenzoic acid (PABA)^12^ and ^18^F-2-PABA^13^, which are involved exclusively in the bacterial folate pathway; ^18^F-maltohexaose^14^ and 6-^18^F-fluoromaltotriose^15^, which specifically target the maltodextrin transporter in bacteria; and ^18^F-fluorodeoxysorbitol (FDS), a synthetic analogue of ^18^F-FDG that selectively localizes in Gram-negative Enterobacteriaceae^16^. However, these radiotracers do not have the ability to distinguish one genus/species/strain from another. Aside from requiring high specificity and selectivity to the target bacterial population, labelling techniques for generating the reporter probes should be simple, rapid and inexpensive to qualify as an ideal imaging agent. Current techniques employ a multi-step process to label metabolites and ligands, which can lead to unnecessary delays and avoidable expenses considering the short decay half-lives (t_1/2_) of some radionuclides.

Our study takes advantage of unique natural molecules that serve as both metal chelators as well as ligands, thus minimizing and simplifying the labelling protocol to just one step. Metal transport is a distinct pathway in bacteria that can be exploited to develop specific probes. Bacteria have developed sophisticated mechanisms for metal acquisition and transport to maintain metal homeostasis within a specified microenvironment. Entire pathways for metal acquisition, comprised of – (i) *de novo* synthesis of metal-chelating “metallophores, Mtps”; and (ii) dedicated membrane transporters that selectively bind the Mtps – have evolved to precisely regulate this process^17^. Mtps are small peptide-like molecules that possess a high affinity for transition metals^18^. The central role that Mtps play as chemical ligands in shuttling metals and the unique biology facilitating this transport into bacteria offers a unique platform that can be harnessed to develop highly versatile and specific contrast agents. Recently, two independent research teams underscored the roles of transport proteins, FyuA and CntA, in copper homeostasis of *Escherichia coli* and *Staphylococcus aureus* respectively. In the *S. aureus* study, a novel metallophore, **<u>st</u>**aphylo**<u>p</u>**ine (StP) (**Fig. 1a**) was shown to chelate metals before the complex is selectively “captured” by the CntA domain and transported into the bacteria by the CntBCDF domains of this ABC transporter protein^19^. On the other hand, pathogenic *E. coli* UTI89 (urinary tract infection isolate) uses the metallophore **<u>y</u>**ersinia**<u>b</u>**ac**<u>t</u>**in (YbT) (**Fig. 1b**) to sequester Cu (II) from the extracellular environment inside the bacteria^20^. Metal bound YbT (**Fig. 1c**) is first selectively “recognized” by its cognate outer membrane protein receptor, FyuA (ferric yersiniabactin uptake A), before the inner membrane ATP-binding cassette transporters, YbTPQ allow cytoplasmic entry. We surmised that these bacterial metal transporters can be targeted by clinically relevant radioisotope labelled metallophores (**Fig. 1d**). Since PET is considered the most sensitive among SPECT, PET, MRI, US and CT imaging technologies^11^, we focused on developing targeted PET imaging contrast agents.

**Figure 1.**
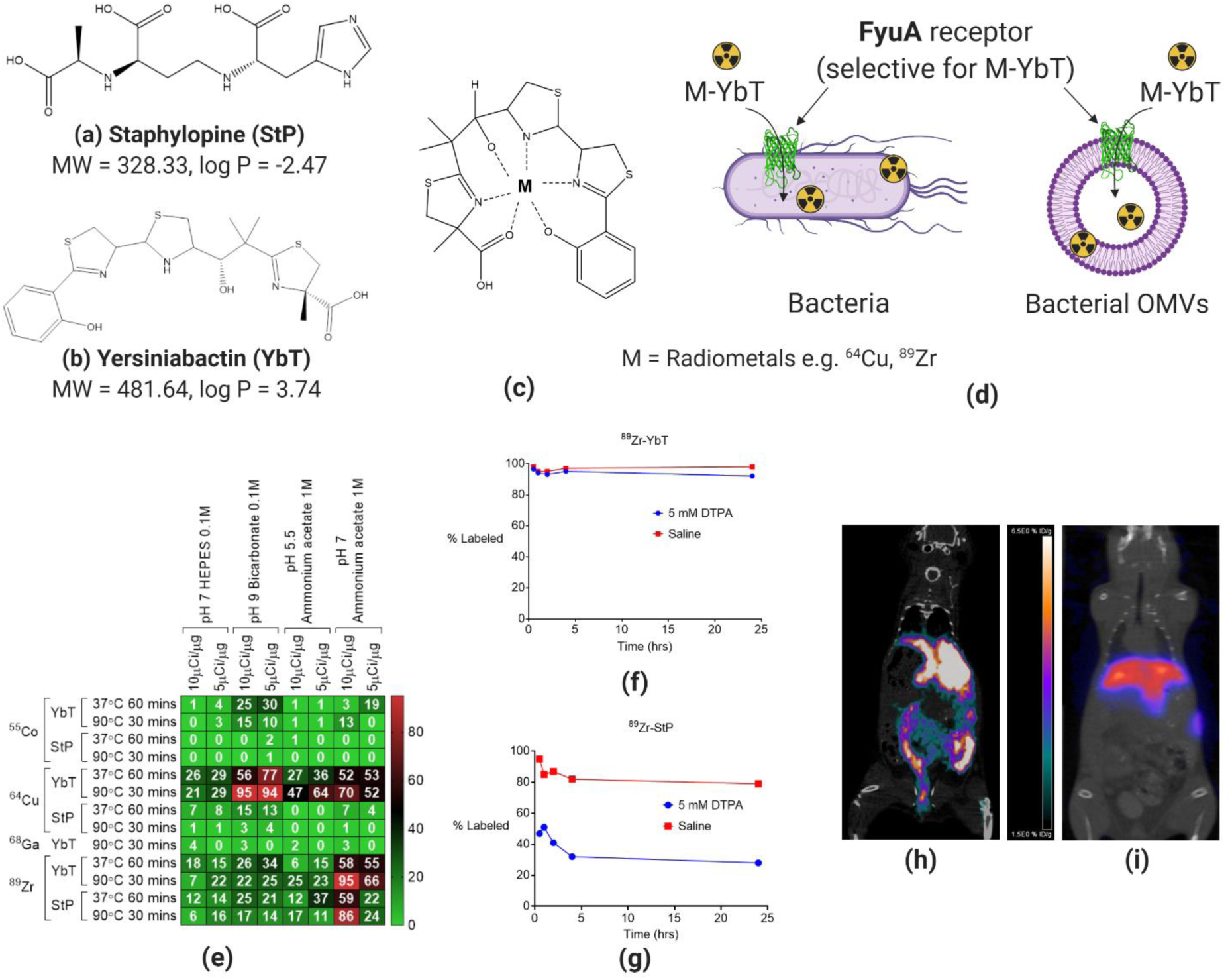
YbT possesses high affinity for ^64^Cu and ^89^Zr and yields stable PET probes. Chemical structures of <u>**(a)**</u> Staphylopine (StP), <u>**(b)**</u> Yersiniabactin (YbT) and <u>**(c)**</u> metal bound YbT. <u>**(d)**</u> Schematic showing selective accumulation of radiolabelled YbT in bacteria and bacterial products. <u>**(e)**</u> StP and YbT complexations with ^55^Co, ^64^Cu, ^68^Ga and ^89^Zr in various physicochemical conditions. Percentage of ^89^Zr labelled <u>**(f)**</u> YbT and <u>**(g)**</u> StP in saline and in DTPA-supplemented saline over 24 hrs. Static PET/CT images of <u>**(h)**</u> ^64^Cu-YbT 24 hrs and <u>**(i)**</u> ^89^Zr-YbT 1 hr post-injection in healthy mice.

## Results

### YbT possesses high affinity for ^64^Cu and ^89^Zr and yields stable PET probes

To optimize the one-step radiolabelling process, an accurate selection of radiometals, Mtps and buffering conditions is crucial for an efficient metal-metallophore complexation. Consequently, we performed *in vitro* radiolabelling of StP and YbT in various buffered and temperature-controlled conditions with four different transition metals – ^55^Co, ^64^Cu, ^68^Ga and ^89^Zr (**Fig. 1e**). We observed that YbT had the highest complexation with both ^64^Cu and ^89^Zr. Since StP manifested moderately high affinity for ^89^Zr, we proceeded to investigate the stability of ^64^Cu and ^89^Zr labelled StP and YbT. When we incubated ^89^Zr-YbT in saline (0.9% NaCl) and in diethylenetriamine pentaacetate (DTPA, a metal-chelator) supplemented saline, ^89^Zr remained bound to YbT in both media for 24 hrs (**Fig. 1f**), but the radiometal dissociated significantly from StP within the first hour in the same conditions (**Fig. 1g**) *in vitro*. We then assessed the stability of YbT probes *in vivo* and observed that both ^64^Cu (**Fig. 1h**) and ^89^Zr (**Fig. 1i**) labelled YbT probes accumulated more significantly in the liver than any other body location, a feature that is likely attributable to the lipophilic nature of YbT.

### ^64^Cu-YbT can selectively identify pathogenic E. coli UTI89 in vivo

Since ^64^Cu and ^89^Zr-labeled YbT proved to have the highest stability amongst all the probes investigated thus far, we wanted to examine whether the YbT probes could selectively differentiate its target, *E. coli* UTI89 (Gram-negative), from representative Gram-negative and Gram-positive bacteria that lack the FyuA receptor *in vivo*. We executed our one-step radiolabelling protocol to demonstrate the ease, simplicity and rapidity of the radiolabelling process from radionuclide acquisition to animal injection (**Extended Data Fig. 1**).

**Extended Data Figure 1.**
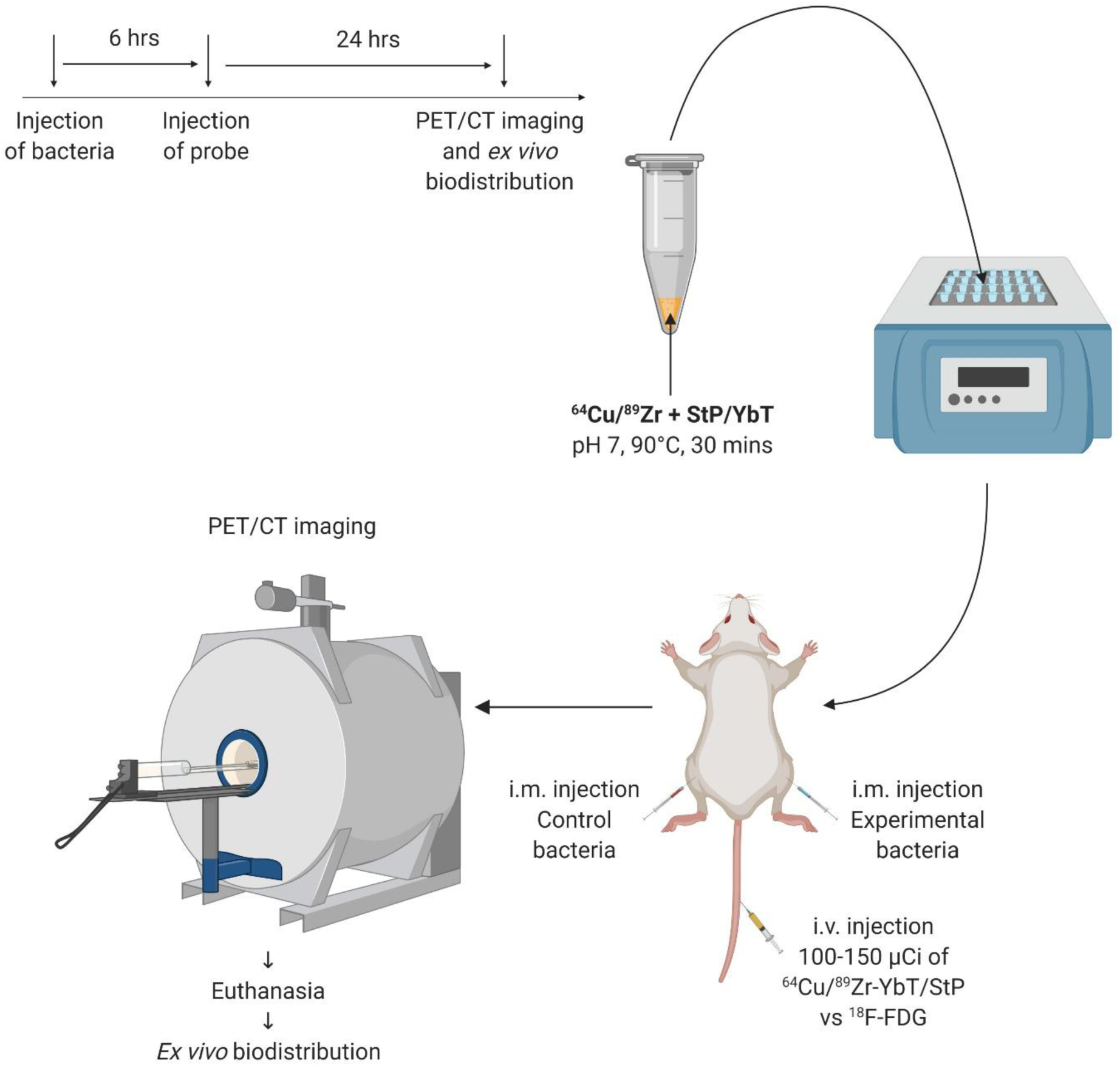
Schematic representing the experimental approach to establish the suitability of radiolabelled metallophores for bacteria-specific imaging.

We selected *Pseudomonas aeruginosa* (Gram-negative) and *S. aureus* (Gram-positive) as control bacteria, since both are widely studied and highly infectious nosocomial bacterial pathogens. Indeed, both *P. aeruginosa* and *S. aureus* lack the cognate FyuA transporters. Tail-vein injections of ^64^Cu-YbT revealed significantly higher accumulation of the probe in UTI89-infected muscles than in *S. aureus* (**Fig. 2a**) or *P. aeruginosa* (**Fig. 2b**) infected muscles, indicating FyuA-mediated *E. coli* species-specific imaging. However, it is likely that there could be a limited degree of crosstalk between strains, particularly within *E. coli* sp. that have the most genetic/pathogenic variants. To determine whether *E. coli* may possess alternative mechanisms to import ^64^Cu-YbT other than the FyuA, we elected to compare the localization of ^64^Cu-YbT in another highly pathogenic strain of *E. coli*, O157:H7 (enterohemorrhagic, EHEC), which does not encode for the FyuA receptor^21^. Intravenous (i.v.) administration of the ^64^Cu-YbT revealed PET signals (**Fig. 2c**), again exclusively in UTI89, thus confirming strain-specific imaging ability of the probe as well. Furthermore, when we administered ^64^Cu-YbT in mice infected with live and heat-killed (90°C for 30 min) UTI89, we observed signals only from muscles harbouring live bacteria (**Fig. 2d**). *Ex vivo* biodistribution revealed concordant results to all the PET/CT images (**Fig. 2e**).

**Figure 2.**
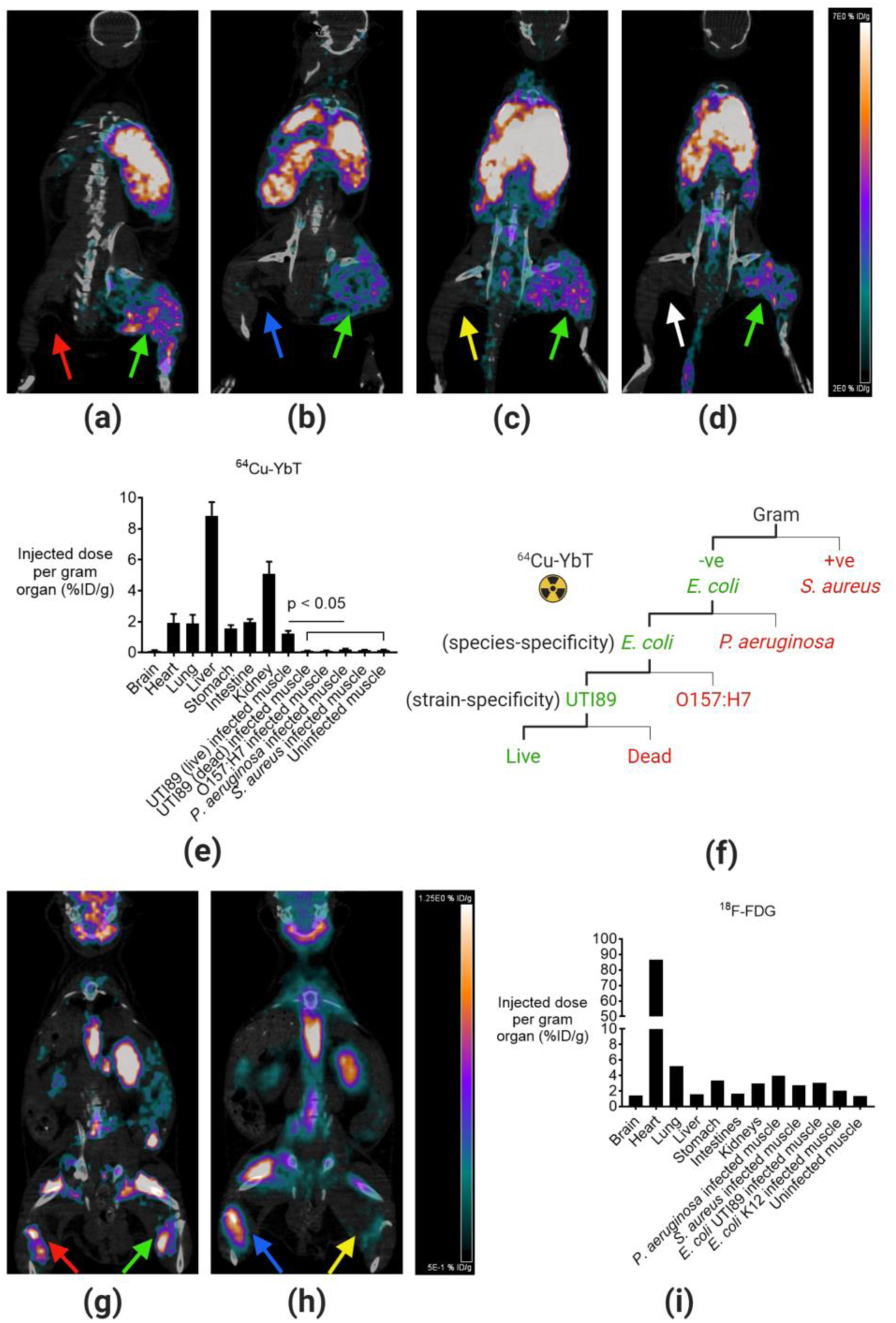
^64^Cu-YbT selectively maps *E. coli* UTI89 *in vivo*. PET/CT images of ^64^Cu-YbT in live UTI89 (green) vs <u>**(a)**</u> *S. aureus* (red), <u>**(b)**</u> *P. aeruginosa* (blue), <u>**(c)**</u> *E. coli* O157:H7 (yellow), and <u>**(d)**</u> dead UTI89 (white) infected mice. <u>**(e)**</u> *Ex vivo* biodistribution of ^64^Cu-YbT 24 hrs post-injection in mice (n=3, results are shown as mean ± s.d.). <u>**(f)**</u> Flow diagram representing Gram, species and strain selectivity of ^64^Cu-YbT in identifying live *E. coli* UTI89 *in vivo*. PET/CT images of ^18^F-FDG 2 hrs post-injection in <u>**(g)**</u> UTI89 (green) vs *S. aureus* (red), <u>**(h)**</u> *P. aeruginosa* (blue) vs *E. coli* MG1655 (yellow) infected mice. <u>**(i)**</u> *Ex vivo* biodistribution of ^18^F-FDG 2 hrs post-injection in mice.

Any new diagnostic probe must be evaluated against a reliable “gold standard”, so we compared the specificity of ^64^Cu-YbT with the clinically available PET probe, **F**luorine-18 labelled **d**eoxy**g**lucose (^18^FDG). We noticed that the latter accumulated in almost equal concentrations in all the pathogens we have investigated to date (**Fig. 2g-i**). It is well-known that ^18^F-FDG is taken up by cells that are involved in both septic and aseptic inflammation^22-24^. Hence, we believe that ^18^F-FDG accumulated within immune cells that infiltrated the infected muscles. The lack of a substantial signal from the muscle injected with the non-pathogenic commensal *E. coli* K12 (**Fig. 2h**) indicates that the bacteria were not able to cause an inflammatory response.

When we attempted to repeat our study with ^89^Zr-YbT, our results showed that the probe failed to selectively identify UTI89 and accumulated mostly in the bones of diseased mice (**Fig. 3a and b**), though previous results indicated sufficient *in vivo* stability of ^89^Zr-YbT in naïve mice (**Fig. 1f and i**). This led us to question whether the intact probe might have dissociated within and eventually effluxed from the bacteria. To confirm this postulate, we performed *in vitro* uptake and retention studies for both the probes. We grew an overnight culture of UTI89 in RPMI supplemented with ^64^Cu-YbT or ^89^Zr-YbT for 2 hrs to mimic the initial bacterial uptake of probes in a physiologically relevant environment. Several studies have shown that bacteria require 2 hrs to accumulate nearly all of the incubated probes^16,20,25^. Subsequently, we pelleted the bacteria and washed with PBS multiple times before we resuspended the pellets in fresh RPMI media for 2 and 24 hrs to investigate how the relative amount of each probe that was retained by the bacteria. **Fig. 3c** shows that while UTI89 was able to retain most of the ^64^Cu-YbT over a period of 24 hrs, the bacteria lost a significant amount of ^89^Zr-YbT over the same period.

**Figure 3.**
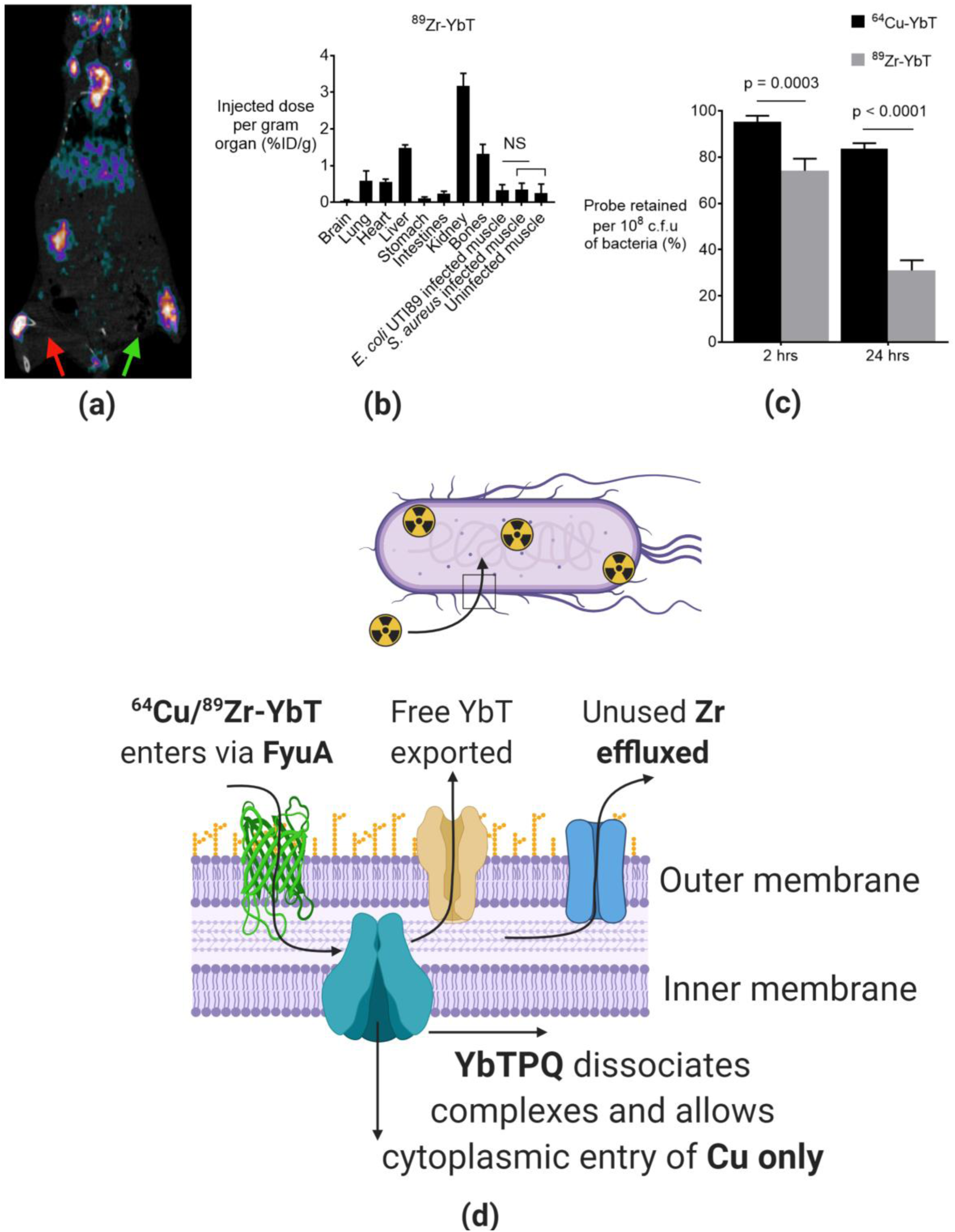
^89^Zr-YbT fails to identify *E. coli* UTI89. <u>**(a)**</u> PET/CT image and <u>**(b)**</u> e*x vivo* biodistribution of ^89^Zr-YbT 24 hrs post-injection in mice (n=3, results are shown as mean ± s.d.). <u>**(c)**</u> UTI89 probe retention 2 and 24 hrs after initial uptake *in vitro* (n=3, results are shown as mean ± s.d.). <u>**(d)**</u> Schematic demonstrating the difference in Cu and Zr processing by UTI89.

### ^64^Cu-YbT specifically identifies FyuA transporter-expressing bacteria

Besides *E. coli* UTI89, probiotic *E. coli* Nissle and pathogenic *Klebsiella pneumoniae* also transport metals using YbT via the FyuA receptor^26-28^. We next reasoned that if radiometal-labelled YbT is truly selective for FyuA only, then our probes should be able to accumulate in Nissle and in *K. pneumoniae* as well. To test this hypothesis, we injected ^64^Cu-YbT in mice that received intranasal administration of PBS (**Fig. 4a**), *P. aeruginosa* (**Fig. 4b**), *E. coli* Nissle (**Fig. 4c**) and *K. pneumoniae* (**Fig. 4d**), with the former two serving as negative controls. The goal was to selectively identify FyuA-expressing bacteria in mice with infected respiratory tracts. After we performed PET/CT imaging and *ex vivo* biodistribution (**Fig. 4e**) of various harvested tissues, we observed significantly higher signals from lungs and trachea of mice that received *K. pneumoniae* and Nissle compared to those that received *P. aeruginosa* and PBS. This illustrates the ability of our probe to trace and identify live bacteria in the pulmonary niche as well.

**Figure 4.**
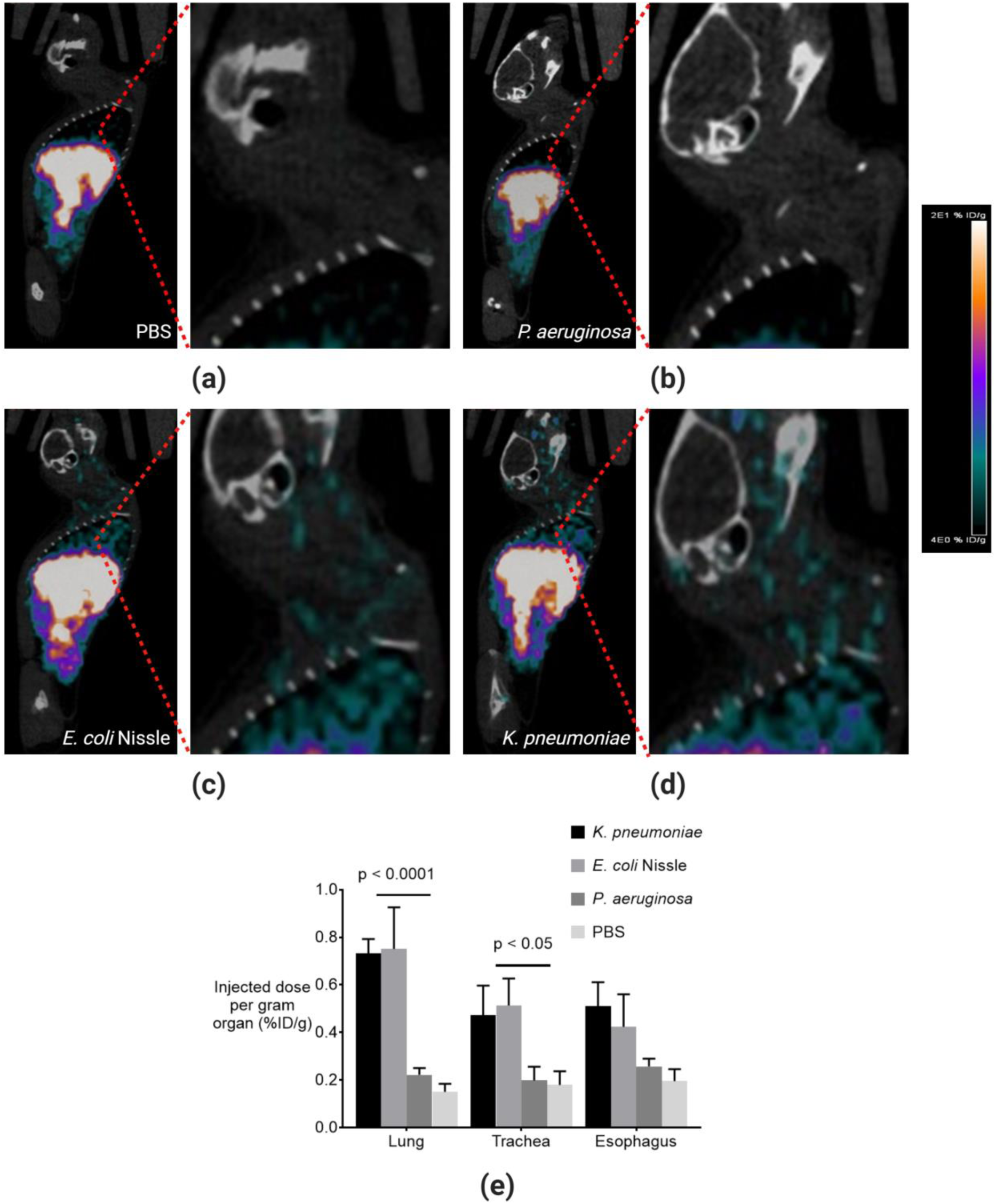
^64^Cu-YbT specifically identifies FyuA-expressing bacteria. PET/CT imaging in <u>**(a)**</u> PBS <u>**(b)**</u> *P. aeruginosa*, <u>**(c)**</u> *E. coli* Nissle, <u>**(d)**</u> *K. pneumoniae* administered mice, and <u>**(e)**</u> *ex vivo* biodistribution of ^64^Cu-YbT 24 hrs post-injection (n=3, results are shown as mean ± s.d.).

### ^64^Cu-YbT can be used to monitor antibiotic treatment efficacy

Clinically, healthcare professionals would prefer to have a “real-time” assessment of therapeutic success or failure for the management of seriously ill patients. To demonstrate whether our probe can potentially be used in such circumstances, we injected ^64^Cu-YbT in mice infected with *E. coli* UTI89 and *K. pneumoniae* to image changes in signal intensity following the administration of the antibiotic, ciprofloxacin. We generated UTI89 clones with a luciferase reporter to track bacterial growth and burden using bioluminescence imaging (BLI). BLI was accurately able to indicate decreases in bacterial burden in response to antibiotic treatment. We used ^64^Cu-YbT to co-register PET signals with BLI, and were able to show a decrease in PET signal corresponding to a proportional decrease in bacterial burden in mice that received two doses of ciprofloxacin (**Fig. 5a**) compared to untreated mice (**Fig. 5b**). When we harvested the thigh muscles from mice post-euthanasia, gamma counter analysis revealed significantly lower radioactive counts from tissues of mice that received ciprofloxacin compared to the control (**Fig. 5c**).

**Figure 5.**
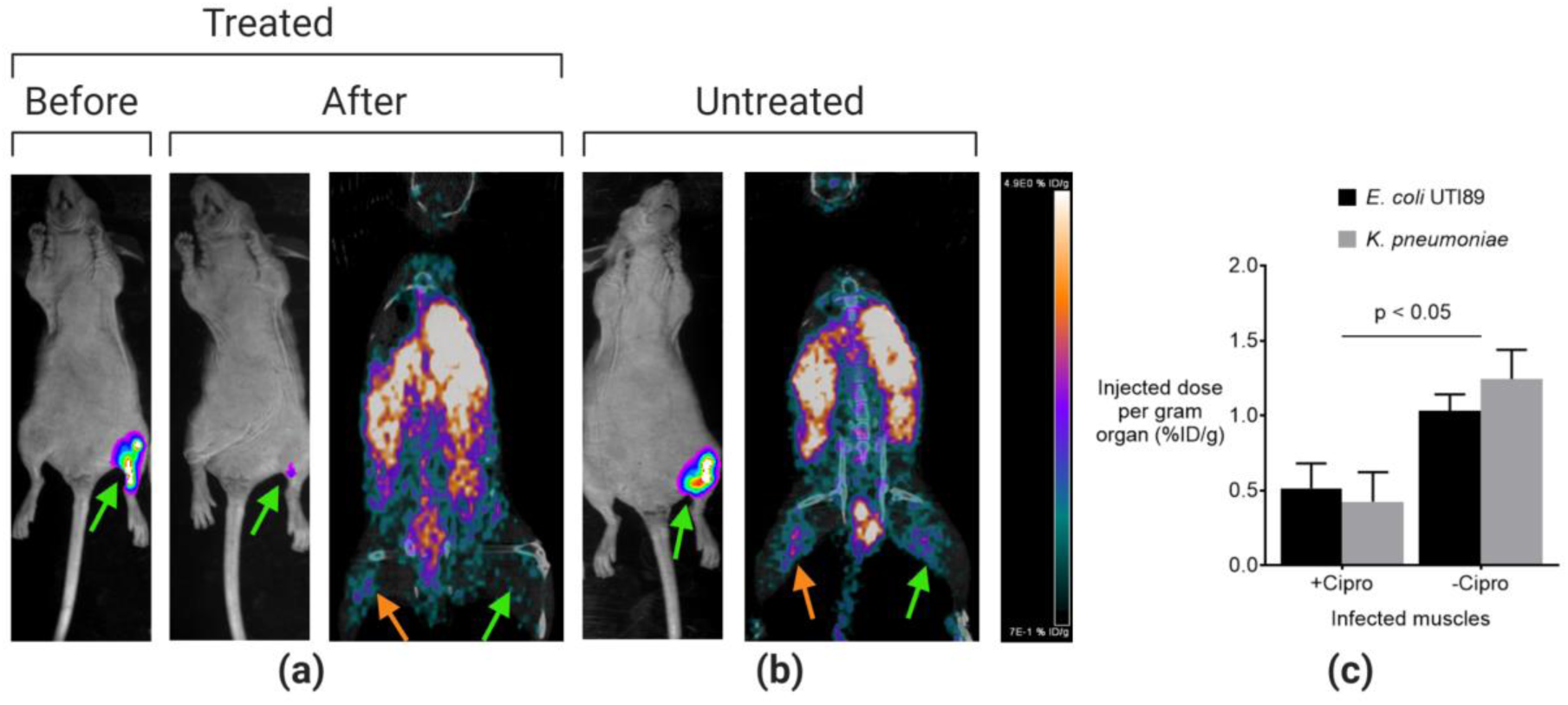
^64^Cu-YbT can be used to monitor antibiotic treatment efficacy. Bioluminescence and PET/CT images of *E. coli* UTI89 (green) and *K. pneumoniae* (orange) infected mice <u>**(a)**</u> treated and <u>**(b)**</u> untreated with ciprofloxacin. <u>**(c)**</u> *Ex vivo* biodistribution of ^64^Cu-YbT in infected muscles of mice 24 hrs post-injection (n=3, results are shown as mean ± s.d.).

### ^64^Cu-YbT can be used to track bioengineered bacteria and bacterial products

The specificity of these distinct bacterial metal transporters for their corresponding metal bound Mtps can be leveraged for *in vivo* visualization of therapeutic bacteria and bacterial virulence factor carrying outer membrane vesicles (OMVs) as well. Since FyuA-expressing *E. coli* species have been proven to naturally possess the metal transporter in their secreted OMVs^29^, we postulated that both Nissle and its secreted OMVs can be selectively tracked by radiolabelled YbT without the need for further genetic manipulation of the bacteria. We tested our theory by injecting Nissle i.v. in subcutaneously developed 4T1 tumour-bearing mice. Usually, greater than 99% of the administered bacteria are cleared from the animals, leaving only a small percentage to colonize the tumor^30,31^. Hence, we allowed 3 days for the bacteria to localize and proliferate in the hypoxic core of the tumour before administering ^64^Cu-YbT in the mice. The control mice did not receive Nissle (**Fig. 6a**). PET/CT and *ex vivo* biodistribution analysis revealed strong signals in tumours of Nissle-administered mice only (**Fig. 6b and c**).

**Figure 6.**
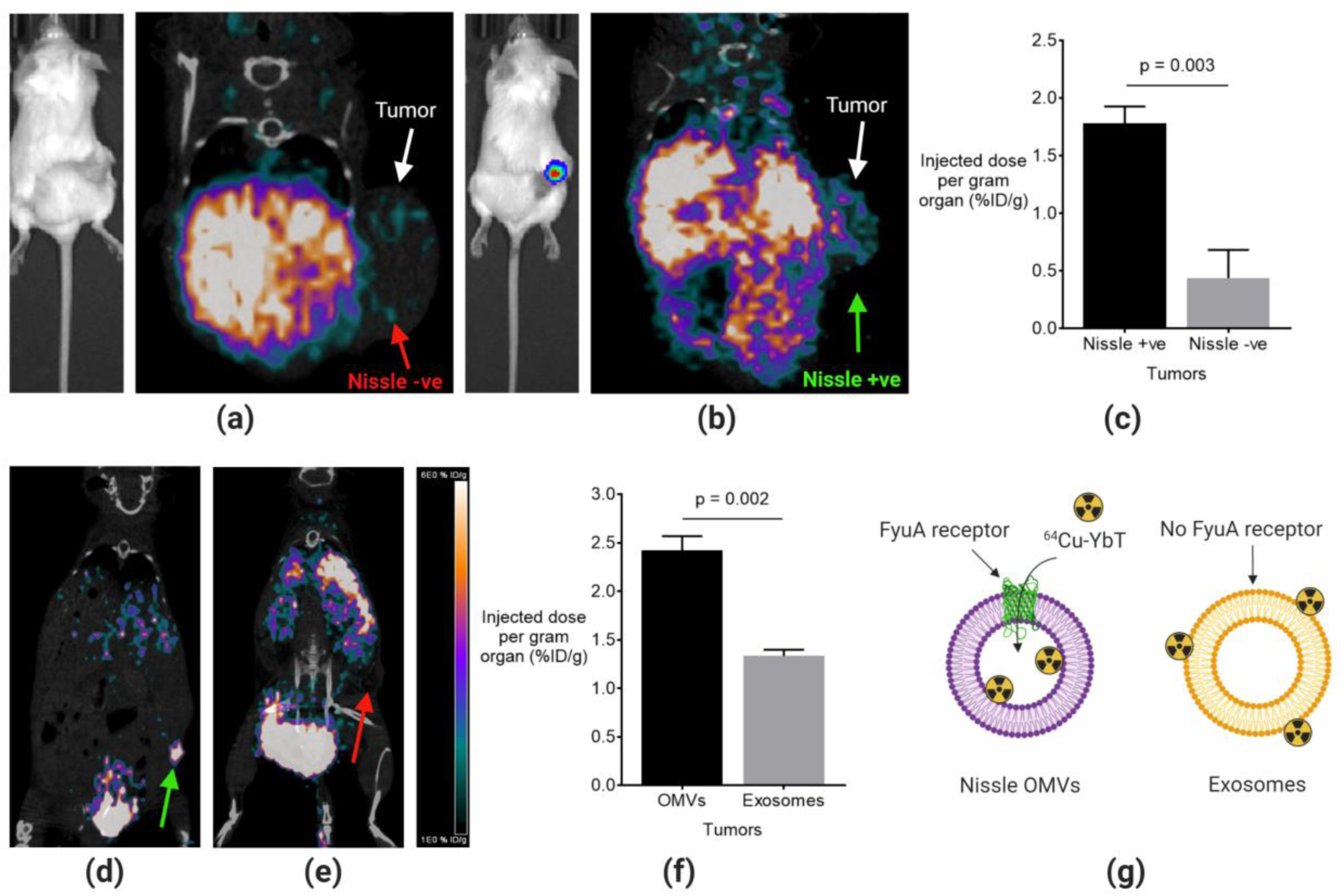
^64^Cu-YbT can be used to track bioengineered bacteria and bacterial products. Bioluminescence and PET/CT images of mice with subcutaneously developed 4T1 tumours <u>**(a)**</u> without (red) and <u>**(b)**</u> with (green) Nissle localization. <u>**(c)**</u> *Ex vivo* biodistribution of ^64^Cu-YbT 24 hrs post-injection in mice (n=3, results are shown as mean ± s.d.). PET/CT images of ^64^Cu-YbT labelled <u>**(d)**</u> OMVs (green) and <u>**(e)**</u> exosomes (red), and <u>**(f)**</u> *ex vivo* biodistribution 4 hrs post administration in mice with 4T1 tumours. (n=3, results are shown as mean ± s.d.). <u>**(g)**</u> Schematic showing transport and trapping of probe inside OMV due to presence of FyuA transporter when compared to nonspecific association of the probes with exosomes that lack cognate transporters.

To achieve our next goal of imaging OMVs in tumours, which accumulate as a result of enhanced permeability and retention (EPR) effect in leaky solid tumours, we first incubated Nissle OMVs and bovine milk exosomes, as the negative control lacking FyuA transporters, with ^64^Cu-YbT to compare the difference in probe uptake between the two types of nanoparticulates. Subsequent purification to eliminate unbound PET probe yielded activities of approximately 200 µCi/ml and 180 µCi/ml for radiolabelled OMVs and exosomes, respectively. After 4 hrs post-administration of the radiolabelled vesicles in our mouse models, we performed PET/CT imaging where we observed radioactive concentration in tumours of the mice injected with OMVs (**Fig. 6d**) compared to those in exosome administered subjects (**Fig. 6e**). We believe that the presence of FyuA on the lipid bilayer of the OMVs allowed selective entry and statistically significant retention of ^64^Cu-YbT within the OMVs (**Fig. 6f**). Since the exosomes lack the metal transporter, ^64^Cu-YbT is merely bound to the outer regions of the nanostructure without being selectively incorporated inside, and eventually had dissociated in the bloodstream *in vivo* (**Fig. 6g**).

## Discussion

In this study, we performed a comprehensive analysis of multiple metal-Mtps and successfully identified YbT complexed to ^64^Cu as a highly stable PET probe that can selectively target bacterial metal transport proteins. Though numerous bacteria-specific PET probes have been developed within the last decade, most of these involve complex and time-consuming reaction mechanisms that eventually yield in tracer synthesis^32,33^. Furthermore, most tracers do not have high complexation (>90%) with the radioisotope, which necessitates an additional purification step before administration. These might prove as potential barriers to clinical translation as hospital/clinical staff members would prefer to be able to prepare the probes in a facile manner. In some of the initial preclinical and clinical investigations, radiolabelled antibiotics such as ^99m^Tc-ciprofloxacin seemed promising, not only for their ability to specifically kill (or disable) bacteria while being nontoxic to human cells, but also for the ease of probe preparation using manufactured kits^34^. However, in later investigations, these SPECT probes proved to not only accumulate in bacterial lesions, but also in sterile inflammatory sites^35^. Metallophore-based probes eliminate the need for sophisticated synthetic chemical reactions since these molecules are readily synthesized and secreted by bacteria, and can easily be obtained using simple purification strategies from culture supernatants^19,20^ (**Extended Data Fig. 2)**. Moreover, we succeeded in optimizing the radiolabelling procedure to a simple and single step of mixing radiometals with these highly selective bioinorganic ligands.

We utilized ^64^Cu-YbT to track pathogenic and probiotic bacteria in three distinct *in vivo* settings – muscle infection, pulmonary infection and intratumorally. *E. coli* UTI89, *E. coli* O157:H7, *E. coli* Nissle, *K. pneumoniae, P. aeruginosa* and *S. aureus* were used for probe testing. The rationale behind selecting the aforementioned bacteria was to demonstrate the highly selective uptake of ^64^Cu-YbT in bacteria only expressing the FyuA transporter protein. In this setting *E. coli* UTI89, *E. coli* Nissle, and *K. pneumoniae* express FyuA, whereas the other organisms represent negative controls. Moreover, we selected these strains to demonstrate differential uptake across bacterial types based upon their phylogenetic hierarchy. For clarity purposes, *E. coli* UTI89, *E. coli* O157:H7, *E. coli* Nissle, *K. pneumoniae*, are Gram-negative bacteria belonging to the *Enterobacteriaceae* family, where *E. coli* UTI89 and *E. coli* O157:H7 are members of the Uropathogenic (UPEC) and enterohemorrhagic (EHEC) subtypes, respectively. In contrast, *P. aeruginosa* is a Gram-negative bacterium belonging to the *Pseudomonadaceae* family while *S. aureus* is a Gram-positive bacterium belonging to the *Staphylococcaceae* family. The central premise of this study was that the probes have minimal cross reactivity and have high selectivity across all major levels – family, genus and serotype – of bacterial class stratification. This is in stark contrast to ^18^FDG, considered a gold standard in clinical infection imaging, which cannot distinguish between different bacteria and essentially is a marker for infection-associated inflammation rather than the presence of “live” bacteria. It is thus critical to ensure that ^64^Cu-YbT was taken up by live and not non-specifically by dead bacteria. Firstly, we displayed how the heat-killed *E. coli* UTI89 never imported our probe. Next, we showed that live bacteria that initially took up the probe, were later neutralized by an antibiotic (ciprofloxacin) regimen and eventually cleared from the infected site. This information could be particularly important in a healthcare setting, where a false positive signal by dead bacteria could mislead physicians to over-prescribe antibiotics in patients, which might eventually lead to the development of multidrug-resistant (MDR) bacteria.

**Extended Data Figure 2.**
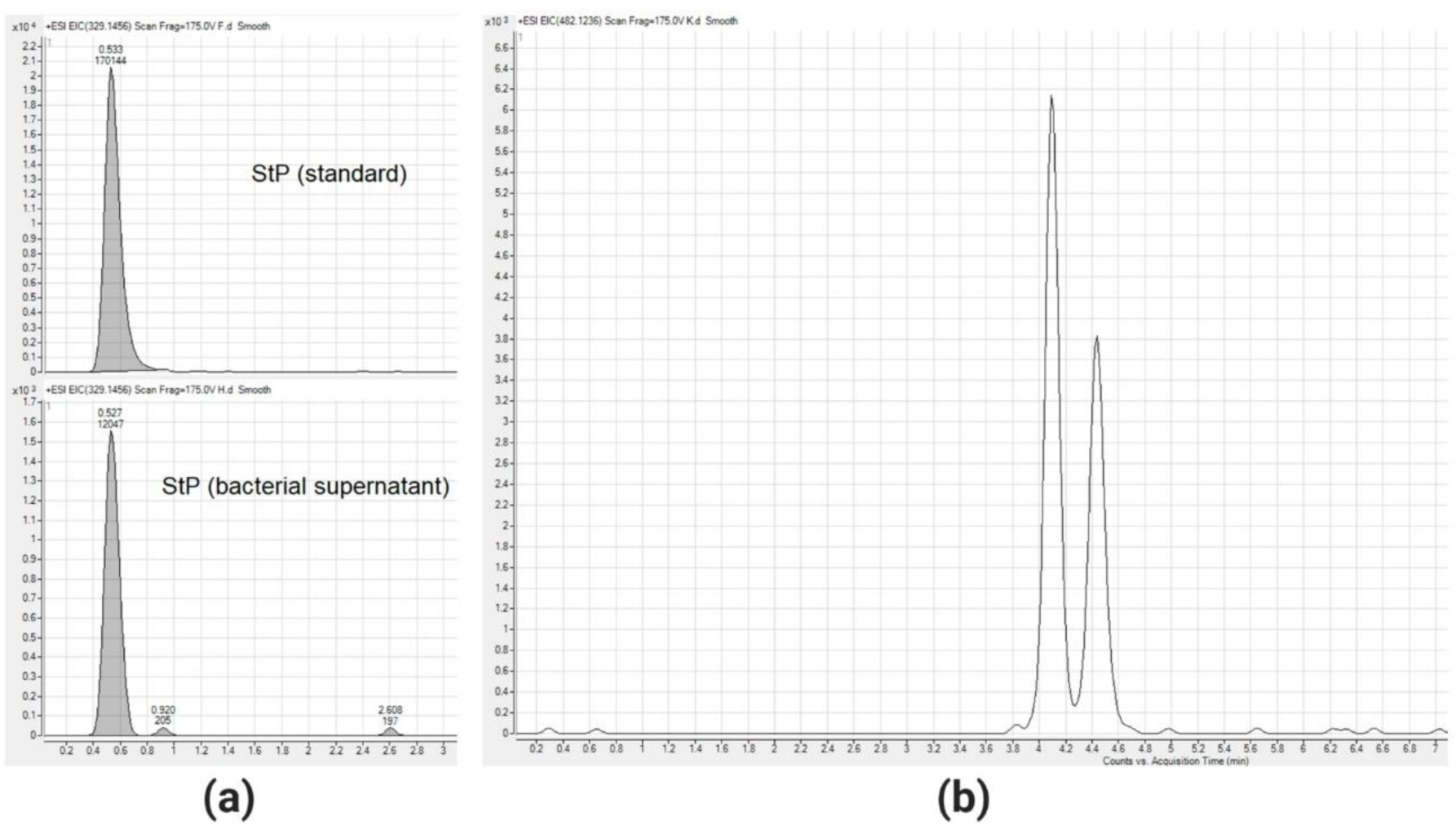
Metallophores are readily available in culture supernatants. M+H^+^ peaks for <u>**(a)**</u> the standard and natural StP molecule synthesized and secreted by *S. aureus*, and <u>**(b)**</u> the two isomers of YbT molecule by *E. coli* UTI89 obtained from LC-MS analyses.

There is precedent for using metallophores as nuclear contrast agents. ^68^Ga coordinated to ferricrocin and desferrioxamine (DFO) has been used in the past to image fungal infections, including *Aspergillus fumigatus*^36,37. 89^Zr coordinated to DFO and coupled to various targeting vectors, including antibodies, peptides and nanoparticles, is a useful strategy for imaging tumor specific receptors. Recently, Pyoverdine-^68^Ga probes were used to selectively image *P. aeruginosa* infections in animal models ^38^. However, due to the short half-life of ^68^Ga (t_1/2_ = 68 min) the ability of ^68^Ga-pyoverdine to track bacteria longitudinally *in vivo* can be challenging. The 12.7-hours half-life of ^64^Cu provides the flexibility to image at both shorter and longer (upto 48h) time scales that allows for optimal clearance of probe and obtaining images with high contrast. Logistically, the longer half-life also allows ^64^Cu-radionuclides to be easily distributed for PET imaging studies at sites remote to the production facility with the loss of approximately one half-life^39^. ^89^Zr has a significantly longer t_1/2_ of 3.3 d which means that some levels of radioactivity can remain in the body for up to a month. This is of particular concern with ^89^Zr-based probes as preclinical studies often report dissociation of the metal from the tracer leading to free ^89^Zr accumulation in the bones^40^. Although ^89^Zr-YbT showed promise in our stability investigations in naïve mice, the bacterial presence in infection models destabilized the probe, which resulted in significant uptake in the bones and joints in subsequent studies. Indeed YbTPQ transporter complex has been shown to be essential for metal-YbT complex dissociation to yield metal-free YbT^20^. Mtps have generally been studied for their affinity for and import of metals such as cobalt, copper, iron, manganese, nickel and zinc, which are deemed “nutritional” by bacteria^19,20,41^. However, zirconium has no known nutritional value to bacteria. Hence it is highly likely that only ^64^Cu was retained inside bacteria and yielded significant PET signals, whereas ^89^Zr was effluxed after 24 hrs from bacteria resulting in loss of PET signal from infection site and subsequent uptake in the bones (**Fig. 3d**). This is interesting because ^89^Zr is known to be a residualizing radionuclide in mammalian cells, which means that upon internalization in cells it is trapped inside the cell^42^. However, since bacterial cells have distinct metal transporters capable of effluxing non-essential metals such as Zr, we suggest ^89^Zr is not a residualizing radionuclide in bacterial cells. Though there have been numerous *in vitro* and *ex vivo* investigations^43-46^, we believe that we are the first to unravel this phenomenon of bacterial metal transport *in vivo* via PET imaging.

While most PET probes have been developed to target *E. coli* and *S. aureus*^*22*^, there has not been any significant effort towards the development of a selective imaging modality for the important pathogen, *K. pneumoniae*. Consequently, we were very keen on understanding how rapidly (**Extended Data Fig. 3**) and selectively ^64^Cu-YbT can provide a positive signal for FyuA-expressing *K. pneumoniae*. While longitudinal imaging would allow physicians to monitor the persistence and/or progression of infection, rapid cross-sectional diagnosis can selectively identify vulnerable patients who require urgent medical attention, particularly in the current and post COVID-19 era. During this pandemic caused by the novel coronavirus, SARS-CoV-2, millions of patients globally have required ventilator-assisted breathing in intensive care units (ICUs). Hospitals in developing countries and make-shift ICUs even in developed countries run the risk of exposing patients to common pathogens such as *K. pneumoniae, P. aeruginosa* and methicillin-resistant *S. aureus* that are known to cause ventilator-associated pneumonia (VAP)^47^. This pathology can be exacerbated by carbapenemase producing *K. pneumoniae* (KPC) and often prove fatal to patients. As scientists begin to unravel the exact pathogenesis of SARS-CoV-2 and emerging respiratory illnesses, early diagnosis of secondary infections caused by bacteria in the respiratory tract would be critical in saving significant numbers of lives globally.

**Extended Data Figure 3.**
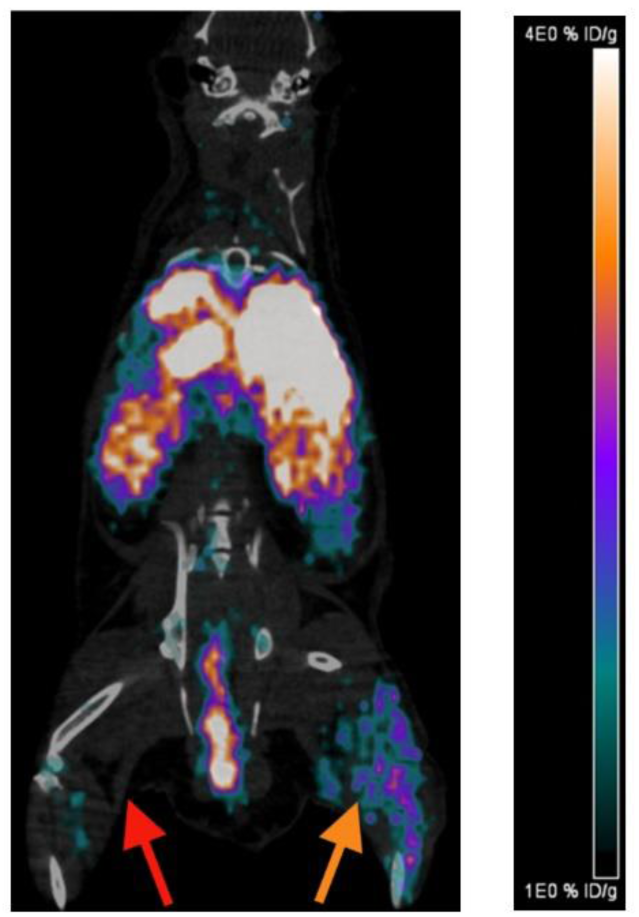
^64^Cu-YbT rapidly identifies *K. pneumoniae in vivo*. PET/CT images of ^64^Cu-Ybt in *K. pneumoniae* (orange) vs *S. aureus* (red) infected mouse 2 hrs post-injection.

We have also validated how ^64^Cu-YbT can potentially be used to advance bacterial therapies using engineered bacteria. Advances in genetic engineering have enabled us to explore the potential of using bacteria, particularly for cancer therapy in the last two decades^31,48-51^. One of the most notable initial studies tested genetically engineered *Salmonella* for its anticancer activity in mouse models of subcutaneously implanted B16F10 melanoma tumors^51^. The results were so promising that a phase II clinical trial was conducted in patients with metastatic melanoma. However, the research ended at that stage because the engineered bacteria did not yield sufficient tumour-targeting efficacy in humans. Selective tracking could have allowed scientists to better map its biodistribution, and hence optimize the therapeutic potential of the engineered *Salmonella*. Whilst numerous optical imaging studies have been conducted, very few nuclear imaging studies have selectively located bacteria in tumors^30^. Optical imaging techniques based on bioluminescence and fluorescence have the inherent drawback of limited light penetration, unlike PET imaging which is depth independent. Recently, tumour localization of OMVs from *E. coli* was confirmed via photoacoustic imaging^48^. While the study described a novel strategy to specifically image the presence of OMVs in the body, it involved an additional step of vesicle design to make it amenable for optoacoustic imaging. Our study illustrates that ^64^Cu-YbT can image bacteria and their products using their endogenous proteins as reporters, without the need for additional genetic modifications. Moreover, the probe can detect bacteria in unique niches, such as tumours, with minimal nonspecific uptake in tumour tissue, thus maximizing the signal to background ratio. Current imaging agents such as ^18^FDG, ^68^Ga-citrate lack specificity as they are unable to distinguish between infection and inflammation^52^. Importantly, imaging agents such as ^18^FDG would not be suitable in this scenario since tumour cells would also uptake the tracer and it would be impossible to delineate signals from bacteria and tumour.

In conclusion, we have demonstrated the successful use of bacterial metallophores as “dual-role” compounds – (i) as a chelator as well as a targeting ligand for imaging and (ii) tracking of pathogenic and commensal bacteria as well as OMVs using a single probe. We have shown the versatility of the probe in detecting bacteria in three different models – myositis, pulmonary infection and an intratumoural niche. Importantly, this technique facilitates precise labelling of “live” bacteria *in vivo*. The small probe size (∼400-500 Da), when compared to conventional antibody or peptide-based PET probes, will potentially allow access through the lining of blood vessels to analyse extravascular structures with relative ease. We anticipate that this will open opportunities to explore the possibility of using the diverse array of natural metallophores, with over 500 examples known to date, as tailored contrast agents for imaging a wide range of wild-type and engineered bacteria. The same probes can be used across diagnostic platforms, for *in vitro* assays using highly sensitive liquid scintillation and gamma counters as well as *in vivo* PET/SPECT/MRI imaging using appropriate radionuclides and metals. For instance, Mtp complexation with radiometals such as ^99m^Tc and ^111^In could facilitate imaging with SPECT scanners that are more ubiquitous and less expensive to operate; and complexation with Mn metal or ^52^Mn radionuclide could facilitate bacterial imaging with MRI or PET/MRI scanners, respectively, to improve spatial resolution. In addition, combining metallophores with novel radiometals can provide insights into unique radiochemistry techniques. Such innovations will reveal new and invaluable information based on metallophore-metal, metallophore-bacteria, metallophore-host and bacteria-host interactions in living systems, which in turn will help design advanced tools and future therapeutic strategies.

## Materials and Methods

### Chemicals

All reagents were purchased from commercial sources as analytical grade and used without further purification. Staphylopine (StP) was purchased from Toronto Research Chemicals Inc. (Toronto, Canada) and Yersiniabactin (YbT) from EMC Microcollections GmbH (Tuebingen, Germany). For animal studies, ^64^Cu and ^89^Zr were obtained from Mallinckrodt Institute of Radiology, Washington University School of Medicine, while ^18^F-FDG was purchased from Cardinal Health (Ohio, USA).

### Bacterial strains and cell culture conditions

Clinical isolates of pathogenic *E. coli* UTI89, *E. coli* O157:H7, *K. pneumoniae, P. aeruginosa, S. aureus*, nonpathogenic *E. coli* K12 (MG1655) and probiotic *E. coli* Nissle were used in this study. Strains were grown in LB agar (BD Difco™) and LB medium (BD Difco™) as appropriate. For bioluminescence imaging, *E. coli* Nissle and UTI89 were transformed with pAKgfplux1, which was a gift from Attila Karsi (Addgene plasmid # 14083; http://n2t.net/addgene:14083; RRID: Addgene_14083) for constitutive expression of lux genes^53^. The transformants were selected on Ampicillin plates prior to use. The murine mammary carcinoma cell line 4T1 (ATCC CRL-2539) was cultured in RPMI containing 10% FCS. The cells were maintained at 37°C with 5% CO_2_ in air, and subcultured twice weekly.

### Metallophore production and identification in culture

*E. coli* UTI89 and *S. aureus* were grown for 20 hrs at 37°C while shaking at 150 rpm. Subsequently, the cultures were centrifuged, and supernatants collected for metallophore identification via liquid chromatography–mass spectrometry (LC–MS). Analyses were conducted using an Agilent 1290 LC-6540 Q-TOF system operated in positive ion mode using an AJS ESI ion source. 5 µL of the samples were injected into an Agilent Eclipse XDB-C18 column (3.5 um, x 100 mm) with a flow rate of 0.5 ml/min and the following solvents: solvent A (0.1% (v/v) formic acid) and solvent B (0.1% (v/v) formic acid in acetonitrile). The gradient program was: 0 – 7 mins 5 – 95% B, 7 – 8 mins 95% B, 8 – 9 mins 95 – 5% B, and 9 – 10 mins 5% B.

### Radiometal-metallophore complexation and stability

^55^Co, ^64^Cu, ^68^Ga and ^89^Zr were produced at the University of Alabama. Complexation was performed by combining 50 µCi of radiometal with 5 or 10 µg of metallophore in 50 µl of various buffered media. Samples are incubated at 37°C or 90°C for 1 hr or 30 mins, respectively. Binding efficiency was determined via iTLC with 50 mM DTPA as the development buffer. The stability of ^89^Zr labelled StP and YbT were assessed at various time points by incubating the probes in saline (0.9% NaCl) and in DTPA-supplemented saline.

### Exosome and OMV preparation

A sequential differential centrifugation protocol was developed to isolate exosomes and OMVs. In brief, high molecular weight (HMW) proteins and fats were removed from raw bovine milk and the supernatant was centrifuged at 91000*g* for 4 hrs to pellet the exosomes. In a similar manner, an overnight culture of *E. coli* Nissle was centrifuged to remove the bacteria followed by concentrating the supernatant using Pierce™ Concentrators (30 kDa, Thermo Scientific™). The concentrated supernatant was then centrifuged at 91000*g* for 4 hrs to pellet the OMVs. Both nanovesicles were suspended in PBS, filtered with 0.22 µm PVDF filters (MilliporeSigma™ Durapore™), and subsequently stored at -80°C until further analysis.

### Radiolabelling for PET/CT imaging

^64^Cu probes were prepared by diluting a stock solution of ^64^CuCl_2_ in 0.1 M sodium bicarbonate buffer to pH 7. A typical reaction involved adding 1 mCi of ^64^Cu to 50 μL of YbT (50 nmol) to bring the reaction volume to 100 μL before incubating on a thermomixer with 800 r.p.m. agitation at 90°C for 30 mins. ^89^Zr probes were prepared by neutralizing 1 mCi (500 µL in 1 M hydrochloric acid) with 500 µL 1 M ammonium acetate (NH_4_OAc) and 2 M sodium hydroxide until the pH was 7. Subsequently, 330 µCi of ^89^ZrCl_4_ (330 µL) was combined with YbT (30 µg in 50 µL 1 M NH_4_OAc, pH 7) and heated at 90°C for 30 mins. Since labelling efficiency for both the metals were > 95%, the probes were administered without further purifications. Exosomes and OMVs were radiolabelled by incubating 1 mL of each of the nanovesicles with 1 mCi of ^64^Cu-YbT for 1 hr at 37°C. Unlabelled radiometal complexes were removed by separating the vesicles with PD-10 columns (GE Healthcare Life Sciences™). Both vesicles were concentrated using Pierce™ Concentrators (30 kDa, Thermo Scientific™) before administration in mice.

### In vitro uptake and retention of ^64^Cu-YbT and ^89^Zr-YbT

Overnight culture of UTI89 was diluted to an OD_600_ of 0.8 in RPMI supplemented with ^64^Cu-YbT or ^89^Zr-YbT (4 µCi each) and grown for 2 hrs while shaking. Bacteria were subsequently pelleted at 8,000 r.p.m for 2 mins and washed with 1× PBS (Sigma) multiple times. Cell-associated ^64^Cu or ^89^Zr levels were measured from pelleted bacteria using a gamma counter. The pellets were resuspended in fresh RPMI media for 2 and 24 hrs after which the centrifugation and washing steps were repeated as above. Experiments were repeated three times from independent bacterial cultures.

### Animal models

6 – 8 weeks old BALB/cJ and athymic nude (NU/J) mice were used for all experiments. For infection studies, 5 × 10^6^ cfu of bacteria were administered either intramuscularly or intranasally 6 hrs or 24 hrs respectively before probe administration. To monitor antibiotic treatment efficacy, ciprofloxacin at a dose of 10mg/kg was given to mice once every 12 hrs via oral gavage. This regimen was thought to provide a plasma peak level in the range that was above those required to achieve efficacy after an oral administration of 500 mg of ciprofloxacin in humans^54^. To image tumour-infiltrating bacteria, 1.5 × 10^6^ 4T1 cells were subcutaneously injected and allowed to develop into a noticeable tumour (size > 200 mm^3^) on the right flank of the mice. This was followed by a single i.v. administration of *E. coli* Nissle (5 × 10^6^ cfu) 3 days before probe PET probe administration. To track OMVs and exosomes in 4T1 tumours, radiolabelled nanovesicles were injected immediately after the tumour reached the appropriate size. All animal experiments were performed under anaesthesia (2% isoflurane) by following protocols approved by the University of Cincinnati Biosafety, Radiation Safety, and Animal Care and Use Committees.

### Imaging and ex vivo biodistribution studies

100 – 150 µCi of PET probes were injected i.v. to selectively image bacteria in both infection and tumour models. 10-20 µCi of radiolabelled vesicles were injected i.v. to study their tumour localization. Small-animal PET scan was performed 4 or 24 hrs post injection on a μPET scanner (Siemens Inveon). Animals were placed in the supine position on the imaging gantry with continued warming for the duration of the scan. A CT scan (80 kVp, 500μA, at 120 projections) was acquired for anatomical reference overlay with PET images for a 15-min acquisition with real-time reconstruction. The PET images were acquired over 15 mins as well and spatial resolution in the entire field of view was determined by ordered subset expectation maximization in 2 dimensions. Histogramming and reconstruction were applied using Siemens Inveon software. Post-processing was carried out with Inveon Research Workplace and general analysis and 3D visualization was used for contouring volume of interest (VOI). These VOI values were considered active infection volumes and used for further analyses.

Bioluminescence images were acquired for 5 mins using an IVIS Imaging System for quantification of radiance (total flux, photons per second, p/s) of the bioluminescent signals from the regions of interest.

After the imaging studies, the mice were euthanized via carbon dioxide inhalation and cervical dislocation. Organs and tissues of interest were removed and weighed. Residual radioactivity in the samples was measured with a gamma counter and results expressed as percentage of injected dose per gram of organ (% ID/g).

### Statistical analyses

All statistical analyses were performed using the GraphPad Prism 7 software. Two-tailed unpaired student’s t-test was performed to compare the means between two groups, whereas one-way ANOVA was used to compare the means among three or more groups. Values of p < 0.05 was considered statistically significant.

## Data availability

The data that support the findings of this study are available from the corresponding author upon request.

## Acknowledgements

We thank Dr. Lisa Lemen and Xiangning Wang for technical assistance with acquisition of all the PET images; Zhiqiang Wang for mass-spectrometry analysis; and Tushar Madaan for bioluminescence imaging of tumor-infiltrating bacteria. We also thank Dr. Pascal Arnoux (Laboratoire de Bioénergétique Cellulaire, Institut de Biosciences et Biotechnology Aix-Marseille (BIAM)) for generously providing us with StP for our preliminary studies. This work was supported by the American Heart Association Innovative Project Award (AHA 18IPA34170157), Department of Defense PRMRP Discovery Award (W81XWH1910125), National Institute Of General Medical Sciences of the National Institutes of Health (R21GM137321) and National Institutes of Health Clinical and Translational Science Award (CTSA) program (5UL1TR001425-04).

## Author Contributions

N.K. conceived the project. N.A.S. and N.K. designed the study. N.A.S. performed all *in vitro* and *in vivo* bacterial experiments. H.A.H. performed radiometal-metallophore complexation and stability studies. S.C.T. prepared exosomes and OMVs. J.R.B. produced ^89^ZrCl_4_ for the study. N.A.S. and N.K. analysed data and wrote the manuscript with input from all authors.

